# Quantifying the Effect of an Acute Stressor in Laying Hens using Thermographic Imaging and Vocalisations

**DOI:** 10.1101/2022.07.31.502171

**Authors:** Huib van den Heuvel, Ali Youssef, Lisette M Grat, Suresh Neethirajan

**Affiliations:** Adaptation Physiology Group, Wageningen University & Research, 6700 AH, Wageningen, The Netherlands; Farmworx, Van der Waalsstraat, 6706 JS, Wageningen, The Netherlands

**Keywords:** Chicken vocalizations, poultry welfare, thermal imaging, laying hens, precision poultry science, Welfare measurement methods

## Abstract

The laying hen sector has multiple issues concerning the animal’s welfare. One crucial factor negatively impacts chicken welfare is stress. The conventional way of measuring and assessing chicken stress is time-consuming and subjective to the assessor. On the other hand, sensing and sensor technologies can be used to obtain objective, continuous and non-invasive/contactless measures of animal behavioural and physiological welfare indicators. The present study aims to investigate the use of thermographic imaging and microphones (sound) in obtaining objective indicators for acute stress in laying hens. During this study, 40 laying hens were stressed by opening an umbrella as a stressor starting from one day age until nine-weeks old. The birds were stressed every other day. Another 12 birds were housed in another cage. These birds were not stressed and served as a control group. The surface temperatures of the bird’s comb and beak decreased (1°C and 2.5°C respectively) in response to the applied stressor. This effect was only seen in the treatment group and not in the control birds. The number of vocalisations the birds produced significantly decreased shortly after stress. The number of calls in the stressed group decreased from 39.5 to 12.1 calls/minute, where the control group decreased from 27.8 to 22.5 calls/minute. It was hypothesized that the number of vocalisations would increase after stress. This difference could be due to the daily behavioural rhythm performed by the birds. The birds might naturally produce more calls at certain hours of the day because of certain behaviours they perform. Three different neural network algorithms were employed to differentiate between the vocalisation of stressed and the control group. This was done by converting the audio files to images and feeding them to the pretrained convolutional neural networks (CNN). The Resnet CNN had the highest categorising accuracy with an overall accuracy of 86 percent. The changes in surface temperature of the beak, comb, eye, and head, as well as the results from the audio analysis could serve as potential indicator for acute stress in laying hens. Future research is warranted to validate the methodologies and findings under different environmental conditions and stressors.

## Introduction

The concern for the welfare of the animals that we keep has risen over the past years. This growing concern comes mainly from a differing outlook on animal ethics (Wise, 2003). However, assessing animal welfare status is difficult because welfare is multifactorial. Furthermore, animal welfare can be measured using both positive and negative indicators. Understanding these aspects of animal welfare is essential in developing and researching the effects of poor or good welfare (Mellor and Beausoleil, 2015).

The laying hen sector has multiple problems concerning the welfare of the animals. A major problem in poultry is feather pecking (FP), which is the act of a bird pecking at or removing the feathers of conspecific birds (Vizzier Thaxton et al., 2016). Such behaviour causes pain and stress in the conspecific animal. While FP is a multifactorial problem and thus can have multiple causations, stress does play a role in a bird becoming a feather pecker (Cronin et al., 2020). Stress can result from multiple sources and can have varied negative effects on poultry. Heat stress in adult laying hens causes a drop in performance and health (Star et al., 2008). Furthermore, chickens subjected to social stressors in the form of unstable hierarchy, produced fewer eggs and performed more agonistic behaviours (Carvalho et al., 2018). This is why, freedom from stress is one of the main points used in the five freedoms assessment (FAWC, 2009).

Measuring stress in chickens can mostly be done by looking at the behavioural and physiological indicators. The problem with conventional techniques of measuring stress is that the results are subjective to the assessor. Therefore, objective measures are needed to have reliable estimate of the stress levels and the response to different stressors in poultry (Banhazi et al., 2012). Physiological measures, such as stress hormones in the blood are of the objective type. However, measuring stress hormones is invasive and time-consuming. The invasiveness of these measures can also stress the birds making the measures unreliable. Non-invasive measurements for stress hormones in chickens exist but are still in development (Weimer et al., 2018).

Precision livestock farming (PLF) is a possible solution to these problems. PLF is the process of automatically recording and managing animals using different sensor technologies. In some animals, such as cows, sensors can be attached to the animal’s body to record its behaviour (e.g., accelerometer). However, this is impractical in chickens due to the size of the available sensors and the number of animals within poultry production systems (Li et al., 2020). Thus, in the poultry sector it is more favourable to use remote and contactless sensors. While most research has been conducted in only experimental settings, some experiments have been done on commercial farms with different sensors (Li et al., 2020). This research focused on using microphones, RFID (Radio Frequency Identification), and image processing to gain information on the birds’ activity, weight, and health.

One sensor that can be promising in birds is the thermographic camera. Many studies have been conducted on using thermography in birds (Moe et al., 2012, 2017; Herborn et al., 2015; McManus et al., 2016; Tabh et al., 2021). The theory behind using infrared thermography (IRT) to detect stress is based on stress-induced hyperthermia and hypothermia. In rats, the core body temperature had risen between 1.5 and 2.0 degrees Celsius after being exposed to a social stressor (Oka, 2018). When an animal is exposed to stress the body (through stress hormones) will drop the body temperature in peripheral areas (such as the comb and wattle in chickens) through the vasoconstriction mechanisms (Moe et al., 2017; Ross et al., 2020). Consequently, the core body temperature will rise due to vasodilation This process is called stress-induced hyperthermia. After the stress event has occurred, the body will try to get rid of the excess heat through vasodilation in the peripheral areas to cool the body back to a thermogenic stable state, which is called stress-induced hypothermia (Moe et al., 2017). Detecting stress using this method has been attempted before with anticipation of feed, handling or change in atmospheric temperature (Giloh et al., 2012; Moe et al., 2012; Herborn et al., 2015; Tabh et al., 2021). However, to what extent minor stressors such as a startle response to a novel object elicited such a response has not been assessed. The current study aims to see whether the surface temperature of the beak, comb, eye, and head can be used as an indicator of stress in laying hens.

Birds are very vocal animals and use their voice for many purposes. The acoustics of birds have been used to detect diseases in animals (Mahdavian et al., 2021). Furthermore, specific types of calls have been used to identify the difference between feather pecking and non-feather pecking flocks (Bright, 2008). A study was also conducted that was able to create an algorithm that could detect the type of stress and its intensity using bioacoustics (Lee et al., 2015). This study aims to see if a simple parameter (the number of vocalizations within set parameters) can be used to give insight in the stress response of laying hens.

## Material and Methods

### Experimental Setup

52 Super Nick chickens were randomly divided into three cages in one stable at the experimental facilities CARUS (Wageningen University and Research). Two cages were 4×4×2m (n=20, treatment cages) and the last cage was 4×2×2m (n=12, control cage). The control cage was enclosed on three sides with cardboard for additional research done on the same animals. The chickens were brought in at three days of age after which they were given a week to acclimatize to their environment. These chicks were vaccinated before arriving as shown in Table 1. The experiment lasted until the chickens were 9 weeks old after which they were sacrificed. The chickens were provided ad libitum feed and water and a perch to sit on, which was introduced after 4 weeks. Air temperature and humidity at the facility changed as the birds aged in accordance with the CARUS protocol for laying hens. Temperature and humidity sensors were hanged on the side of the control cage and in-between the treatment cages. Daily temperature and humidity were recorded to correct for the change in ambient air temperature (Appendix 1). The birds were kept at a day-night cycle of 16:8 hours, from 06:00 till 20:00. The chicks were exposed to different types of light-emitting diode (LED) lights as part of another ongoing research. Only the treatment cages were exposed to these LED lights. The birds in the control cage were always exposed to the same white light. During the measurement days of the present thesis, no light-colour manipulation was performed (Appendix 2).

**Table 1.** The vaccinations the chicks received at day 1 of age

| **Vaccination** | **Disease** |
| --- | --- |
| **Rispen** | marek |
| **Novamune** | *infectious bronchitis* |
| **Avinew** | *newcastle disease* |
| **Evalon** | *coccidiosis* |

To induce stress in the birds an umbrella was used. The umbrella was opened in front of the birds to startle the birds and cause a flight response. Only the birds in the treatment cages were stressed during the stressing events. Measurements were done on 14, 16, 18, 22, 24, 28, 30, 32, 36, 38, 42, 44, 46, 50, 52, 56, 58, 60 and 64 days of age (Appendix 2).

### Thermography

To investigate the possibility of using thermography to assess the stress response of chickens a Flir1020 thermal camera was used (1024×768 resolution, + 1°C accuracy, emissivity 0.95). Thermographic images were taken at three different moments: an hour before, just after and an hour after inducing the stress response. Per moment 40 pictures were taken of either the left or right side of the chickens’ face (fifteen in both large cages and ten in the small one). The pictures were taken between 0.5 and 1 meters away from the bird. For this study, the chicks were not marked and thus it was not possible to identify individuals. Therefore, when taking a picture, chicks were chosen at random. No pictures were taken of the same chick twice during the same measuring moment. Furthermore, no thermographic pictures were taken of any birds after they drank water or those with a soiled beak.

The thermographic images were analysed using FlirResearch software. During the analysis regions of interest were drawn in the program and the maximum, minimum and average temperatures within these regions of interest were extracted (Figure 1). The selected regions of interest were the beak (from the nostril to the tip of the beak back to the base of the beak), the eye (an oval in around the inner eye), the comb (from the nostril up to the end of the comb excluding the parts sticking out at the top or hair covering at the base), and the head (an oval circling the head from the beak up to the comb around the ear canal and back to the beak).

**Figure 1.**
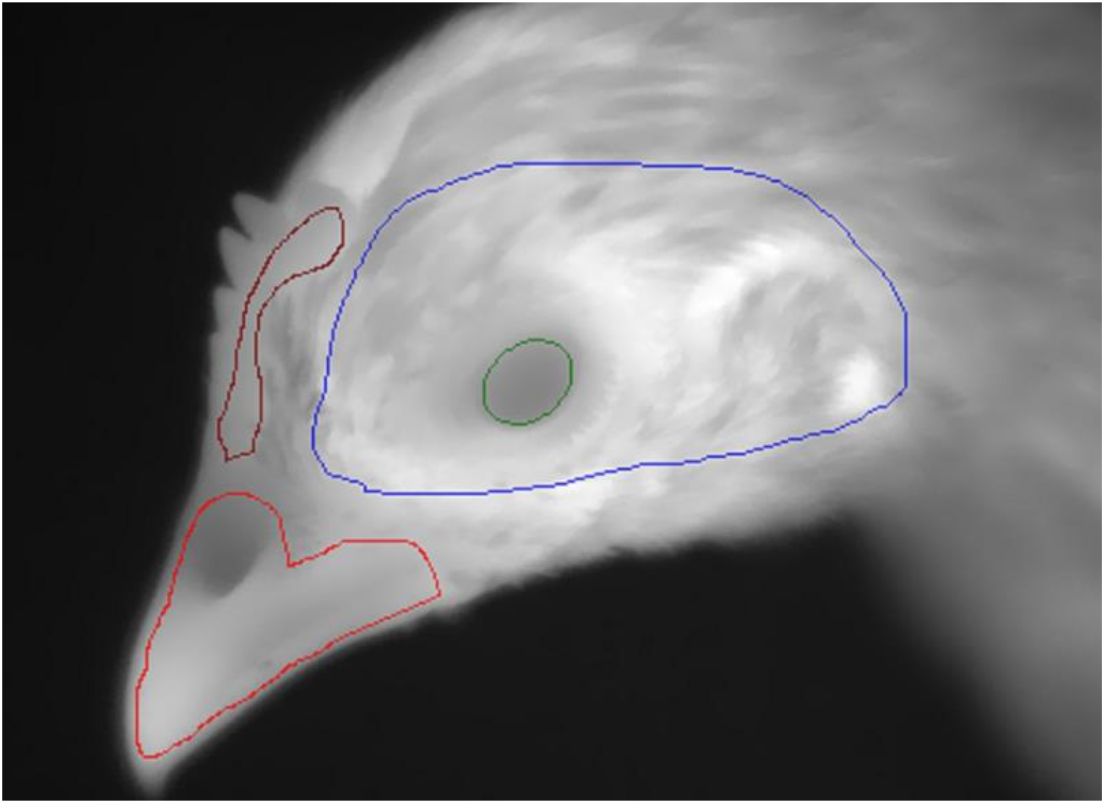
Thermal image of closeup view of the laying hen from this study. The regions of interest used in the analysis are shown in circles. The red color circle represents the beak. The brown circle represents the comb region of the body. The green color circle represents the eye. The blue circle represents the head

To assess the effect on temperature in the regions of interest (beak, eye, head and comb) during stress a general linear model was performed in IBM SPSS statistics 27.

The model had the time of measuring (during, before and after stress) and whether the birds where from the control or treatment (stressed) group as main effects. Furthermore, the interaction between the treatment and time of measuring was included in the model. The cage’s ambient temperature and the cage the birds were in (1,2 or 3) were included in the model as covariances. In total 2280 photos were analysed (Table 2). However, since the comb of the birds did not develop until a later age this region of interest could not always be analysed. Since the ambient temperature changed as the chicks were growing up (as per protocol) the age of the birds could not be used in assessing the effects of stress on the temperature of the different regions of interest. The data was checked for a normal distribution on the residuals. Normality was assumed when the kurtosis and skewness were within −2 and 2.

**Table 2.**
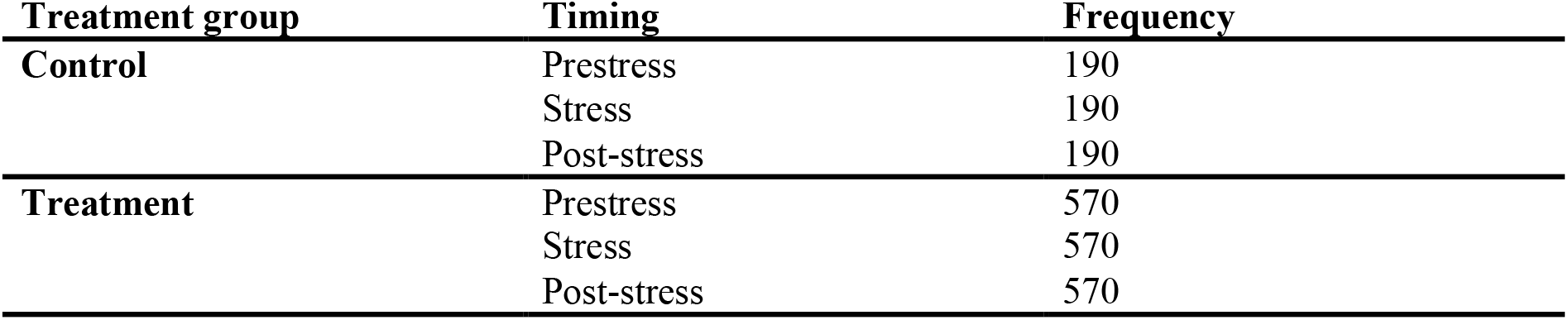
Frequency table of the thermographic pictures

### Audio signal analysis

Recordings were taken from the sounds in the cages for one hour before and one hour after stress. During the first three weeks, these measurements were taken from the second and third cage (both treatment cages). After week three, they were taken from the control (cage 1) and treatment cage (cage 3). Two microphones were used in this study. The first is a Tascam microphone, and the second is a Rode Video Micro which was connected to a GoPro video camera. Both microphones were suspended two meters high in the middle of the cage. In total 42 hours of audio data was collected, and two approaches were taken in this study. First approach looked at single call expressions from the birds, and the second looked at determining group level audio vocalizations signals which are the current practical reality in rearing the birds in commercial production floor.

### Variety in the Amount of Calls

After collecting the audio files, the hour-long recordings were cut with Audacity© (Audacity team, 2022). For the post-stress condition, the first 10 minutes were used for analysis. For the prestress condition, the time between 10-20 minutes and 40-50 minutes were averaged and used for the analysis. The cut files were analysed using a customised MATLAB-based sound processing algorithm. The algorithm extracts and counts the number of vocalisation events based on predefined energy and frequency ranges. The energy and frequency ranges is defined by specific hyper-parameters for the algorithm. These parameters are determined based on visual investigation and trial and error approach. The defined parameters are the frequency of the sound (3-4.5 kHz), the duration of the call (2+ 0.5 seconds) and the minimal energy of the call (500). The number of occurrences within the file was counted and converted to number of calls per minute per cage.

From day 28 onwards, the data were complete for the control cage and one treatment cage. Therefore, the number of calls was analysed with a general linear model in which treatment (control or stressed birds) and the stress condition (prestress and post-stress) and their interaction were included as explanatory variables. The age of the birds in days was included as a covariate in the analysis. The data was checked for a normal distribution on the residuals. Normality was assumed when the kurtosis and skewness were within −2 and 2.

### Recognition of stress calls

Convolutional Neural Networks (CNN) are feature extractors and are deployed for image analysis. CNN are deep forward neural networks and made of interconnected neurons that have inputs with biases, learnable weights, and activation functions. In this study, audio signals were converted to images and the CNN were used for extracting information regarding the image (Figure 2). Upon extracting the features, the CNN Models helped as a preprocessor of the data before feeding it to the network architecture. The network consisted of three layesr as follows, a convolutional layer, pooling layers and connected layers (Singh and Manure, 2019). In this study, 3 types of networks were used for the audio data analysis namely ResNet, AlexNet and GoogleNet.

**Figure 2.**
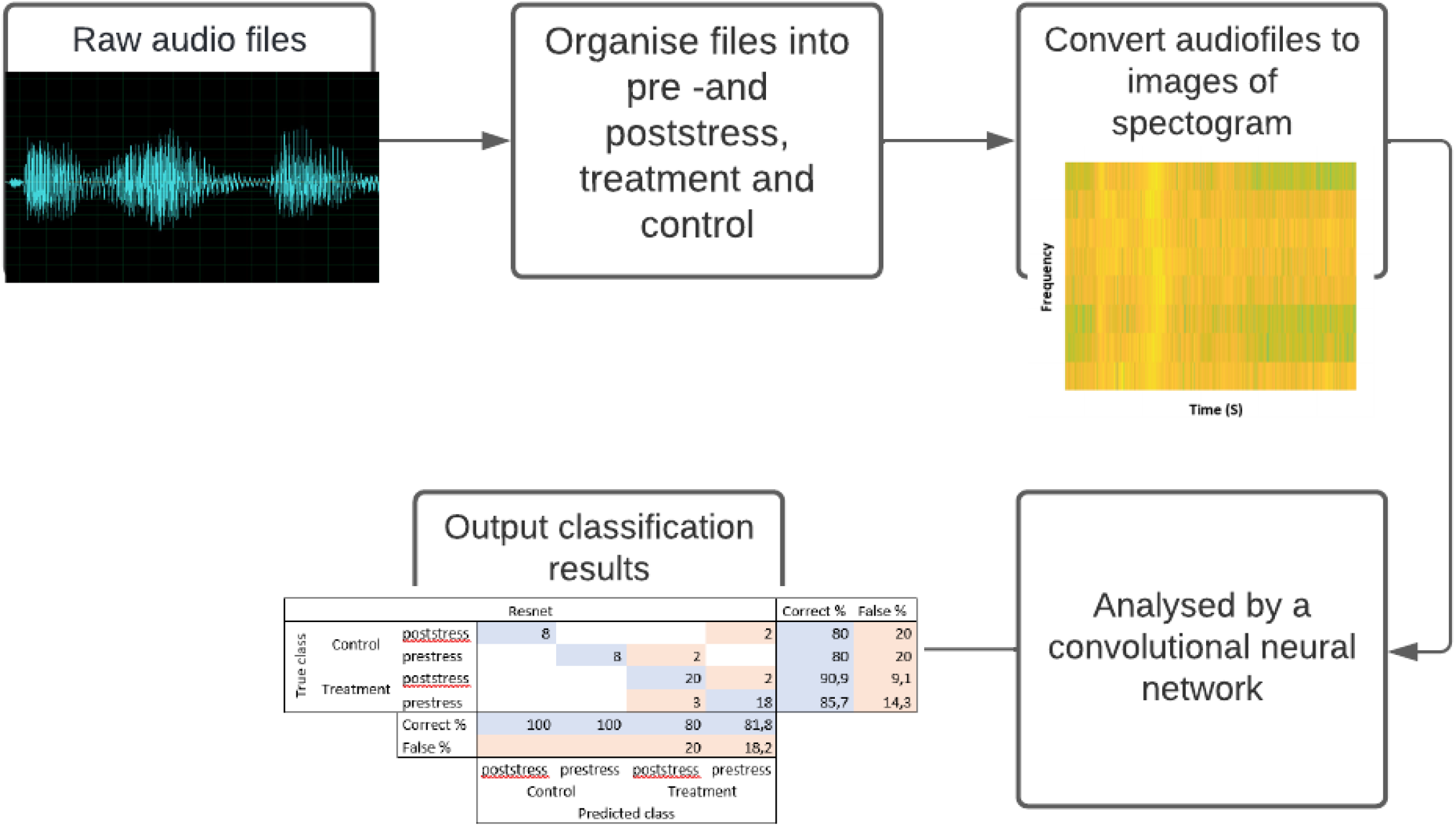
Flow diagram visualizing the process of classifying audio files using convolutional neural networks

AlexNet CNN is the winner of the ILSVRC competition in 2012 and is made of connected and stacked layers and includes 5 convolutional layers followed by 3 fully connected layers and max-Pool layers in the middle in the form of a sandwich. For achieving faster training and classification, AlexNet uses rectified linear unit non-linearity process (Krizhevsky et al., 2012; Iandola et al., 2016).

GoogleNet CNN is the winner of the ImageNet ILSVRC competition in 2014 and introduces a novel ‘Inception’ module and is made of 22 layers for training and 5 POOL layers (Szegedy et al., 2015). This is composed of a subnetwork of parallel convolutional filters and their outputs are concatenated.

Resnet is the winner of the ILSVRC 2015 challenge award and used a 152 layered (He et al., 2016), deep CNN. The niche of ResNet in comparison to other deep CNN, is that it uses shortcut connections and skip connections (network-in-network architecture). To simply put, the residual learning process was shortened by allowing the input data to ‘bypass’ through the layers. ResNet offers a faster and more efficient way to solve problems the deep neural network faces (Glorot and Bengio, 2010; Khan et al., 2020).

These three pre trained neural networks were provided the collected audio data from this study converted to spectrograms. These spectrograms were analysed by the CNNs and categorised into either prestress or post-stress and control or treatment. The CNNs learned how to categorise these images into the correct category over time based on differences in the spectrogram. The accuracy with which it was able to categorise these files was used as an indicator of its use in recognizing stress calls in laying hens.

## Results

### Thermographic imaging

The correlation between the maximum ambient temperature and the average temperature of all the regions of interests was found to be significant (p<0.001)

Right after applying the stressor, the average temperature in all investigated body areas declined significantly (p=0.00) when compared to the prestress and post-stress condition. No significant differences in surface temperature were found between the prestress and the post-stress conditions. Furthermore, it was found that the maximum daily air temperature had a significant (p=0.00) influence on the average surface temperatures of all regions of interest.

This change in temperature was different in intensity for the different regions of interest. In the beak area the average temperature decreased by 2.5°C when the chicken was exposed to stress (Figure 3). When exposed to stress, the eye temperature decreased by 1.0°C (Figure 4). The temperature in the head region decreased the least with a 0.8°C (Figure 5). The average comb temperature decreased the most during stress with a 3.4°C degree drop (Figure 6). After an hour the temperature in all regions did not differ from the prestress temperature.

**Figure 3.**
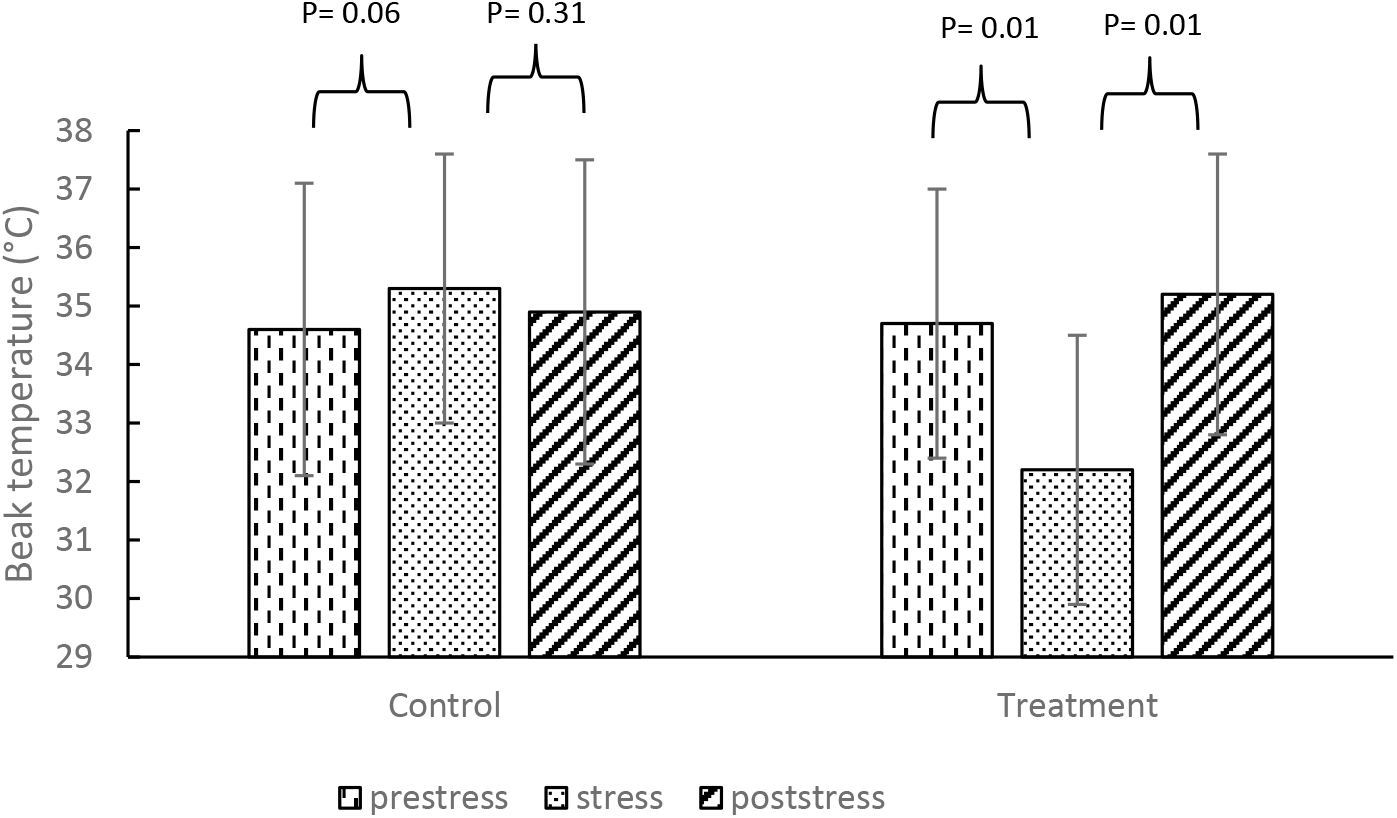
The average beak temperature (°C) before, during and after stress. The results are split between the control and treatment group. The error bars represent the standard deviation (n=2258)

**Figure 4.**
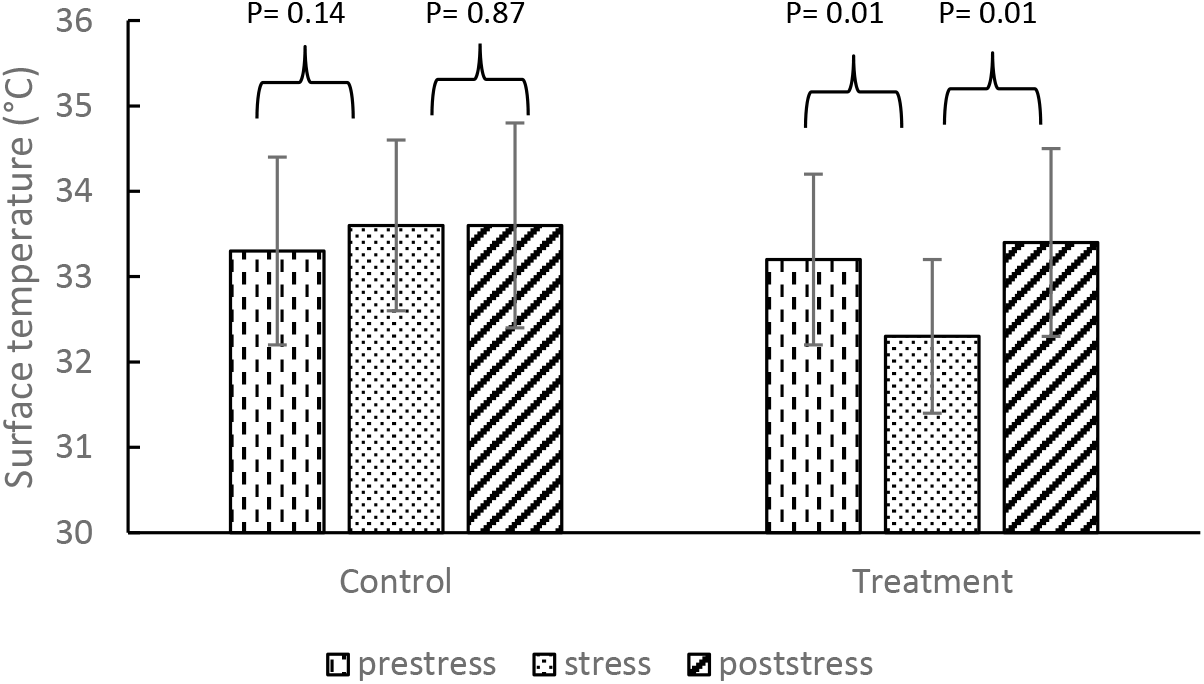
The average eye temperature (°C) before, during and after stress. The results are split between the control and treatment group. The error bars represent the standard deviation (n=2254)

**Figure 5.**
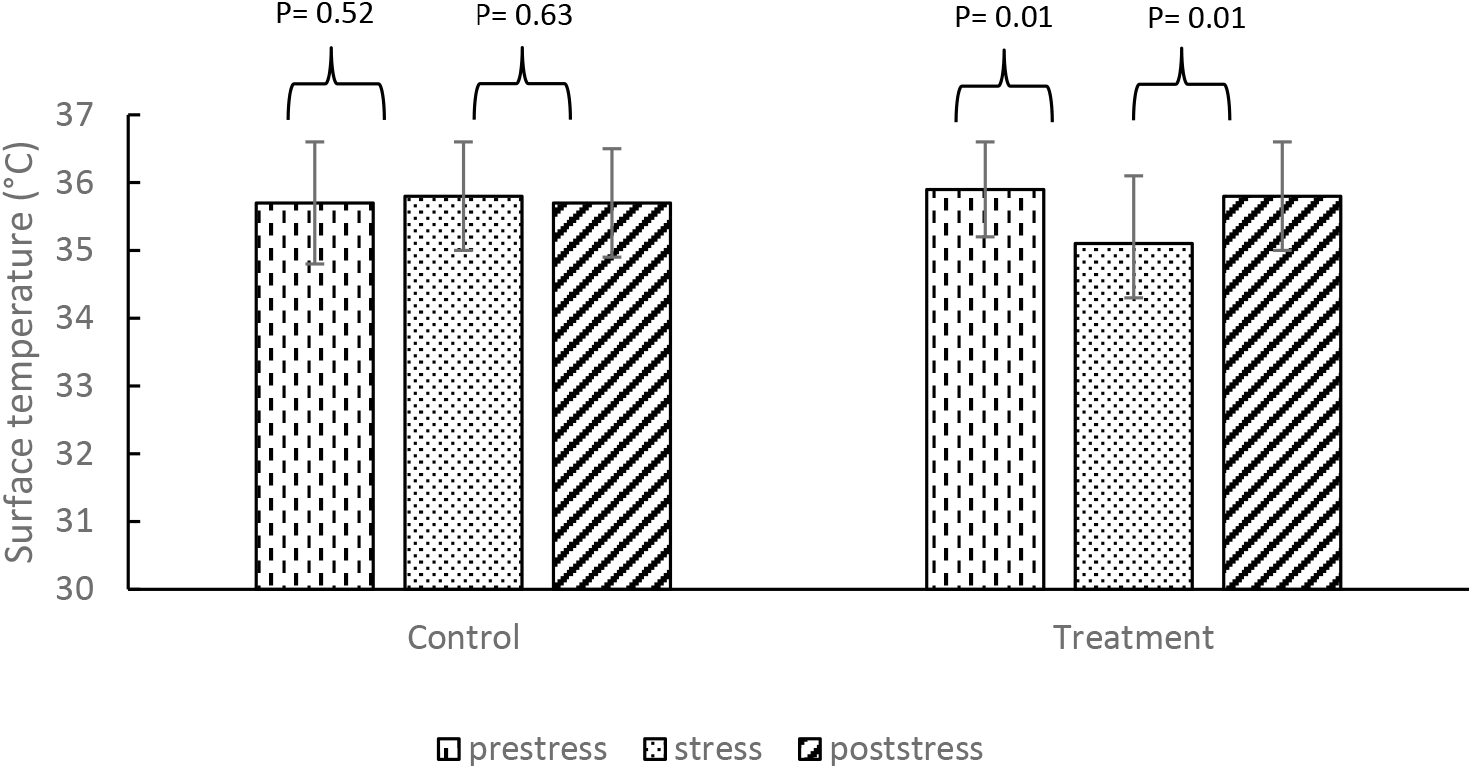
The average head temperature (°C) before, during and after stress. The results are split between the control and treatment group. The error bars represent the standard deviation (n=2264)

**Figure 6.**
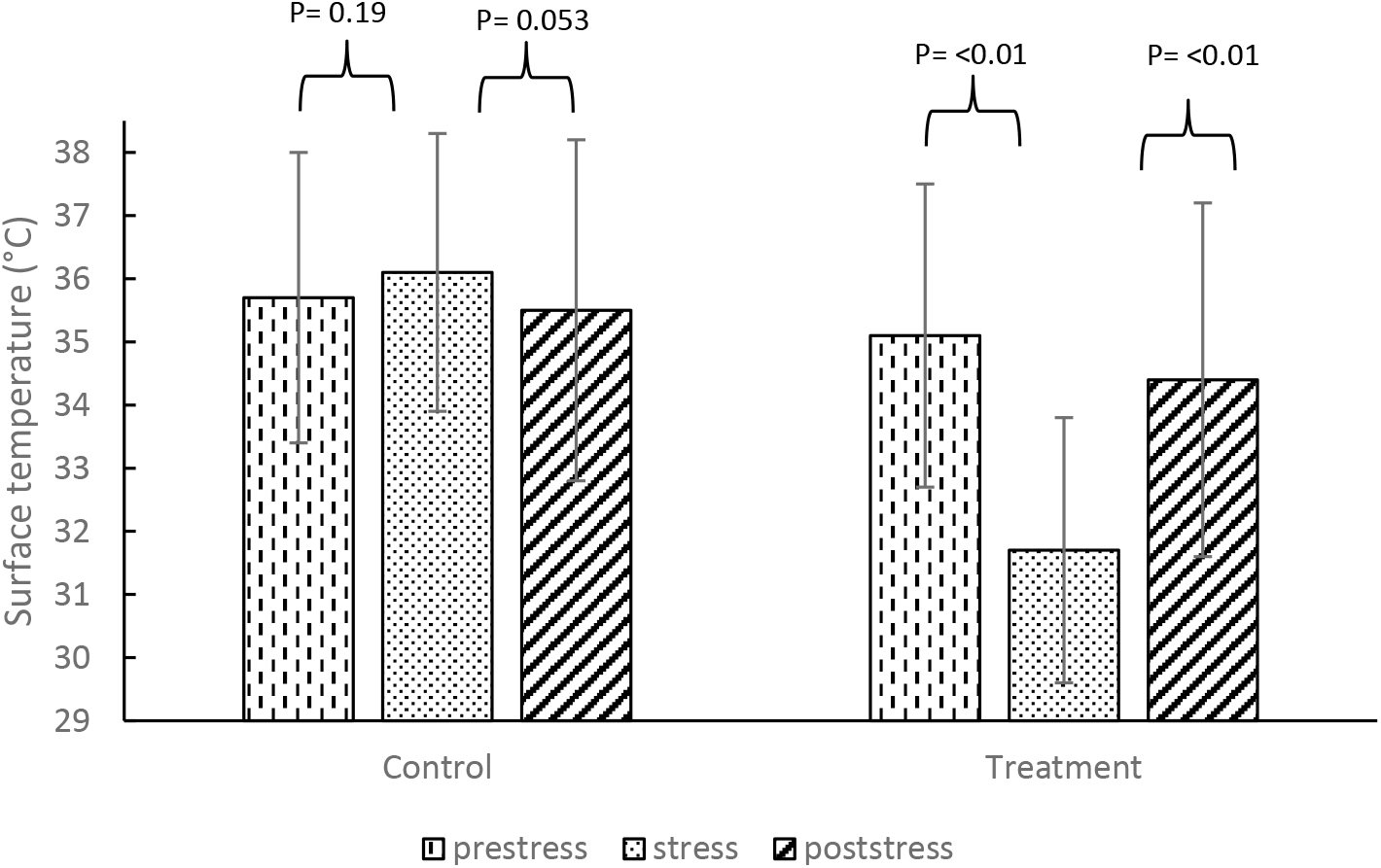
The average comb temperature (°C) before, during and after stress. The results are split between the control and treatment group. The error bars represent the standard deviation (n=995)

### Sound analysis

#### Variety in the number of calls

The interaction between treatment and timing was significant(p=0.021). Only in the treated (stressed) group the number of calls was significantly lower after stress compared to prestress, the calls decreased from 39.5 to 12.1 calls/minute (p=0.001), while this was not significant in the control group (27.8 to 22.5 calls/minute; p=0.85). No significant effect of age on the number of vocalisations was found (p=0.221).

#### Recognition of stress calls

After 300 iterations of the three CNNs, the Resnet gave the best classification (Appendix III) result with 85.7% validation accuracy (Figure 8) compared to 55.6% for Alexnet (Figure 9) and 74.6% for GoogleNET (Figure 10).

**Figure 7.**
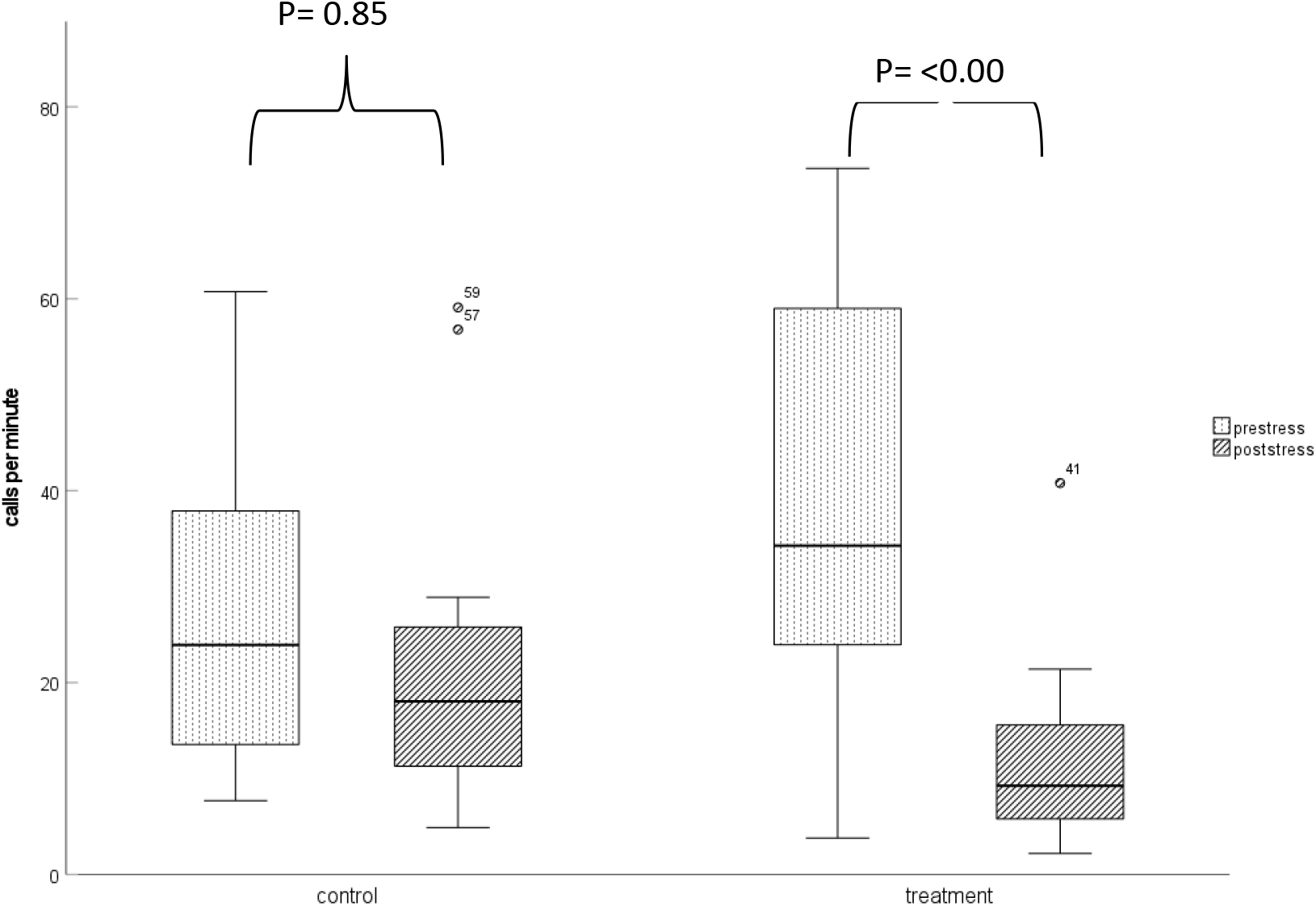
The average calls per minute produced by the control and treatment (stressed) chicks before and after the stressor was applied stress. The dots represent outliers in the analysis. The whiskers represent the minimum and maximum and the middle line

**Figure 8.**
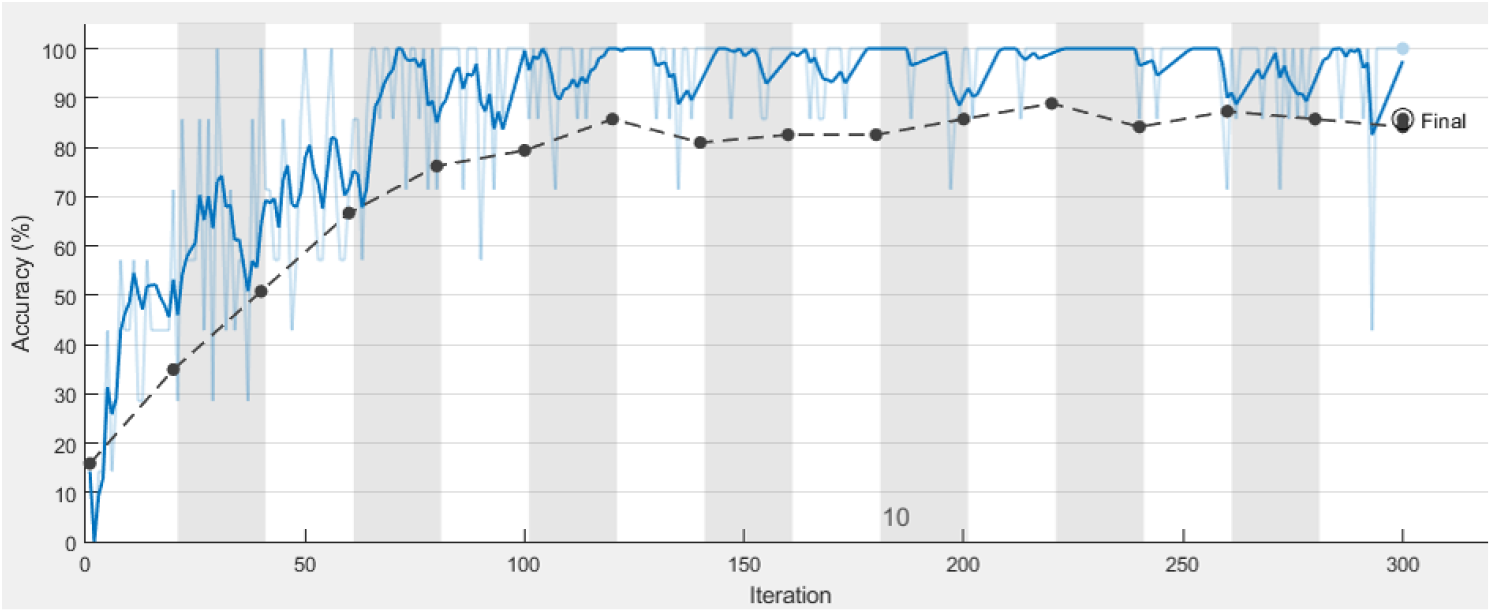
The accuracy of the Resnet CNN over the iterations during training. The dark blue line represents a smoothed version of the training of the CNN, while the light blue line represents the accrual result from the training. The black striped line represents the accuracy results from the validation after 20 iterations.

**Figure 9.**
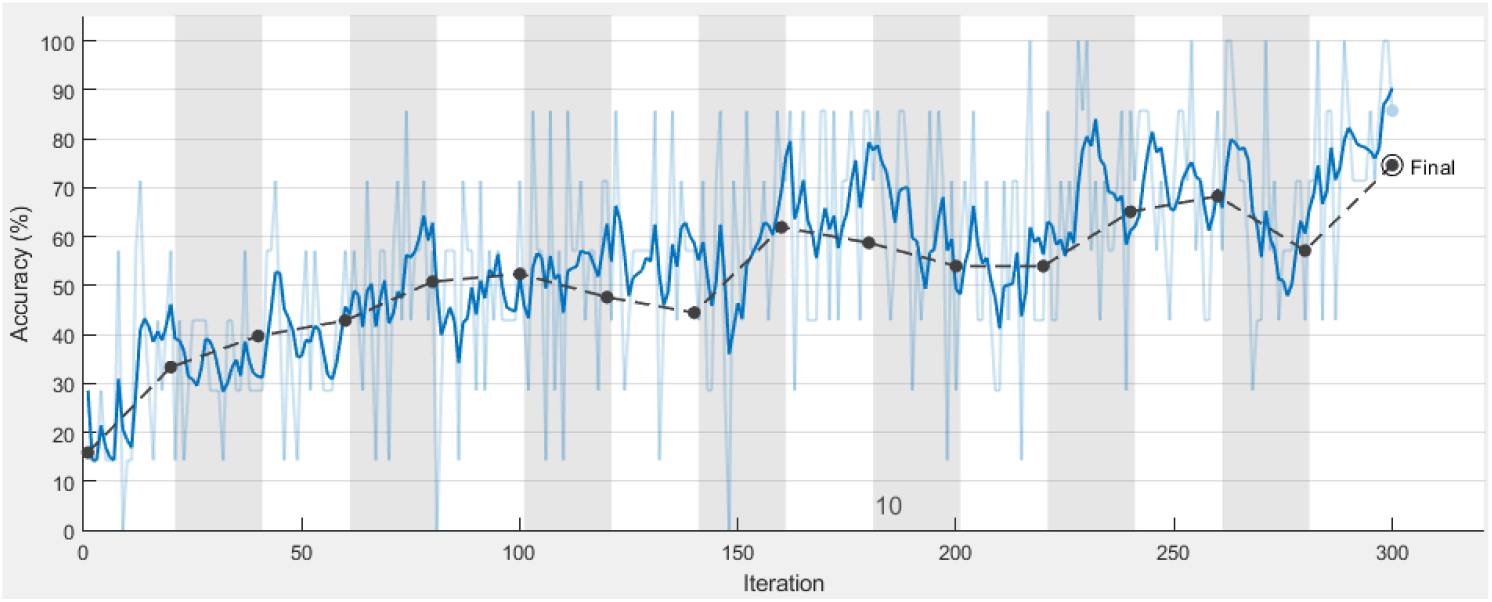
The accuracy of the Alexnet CNN over the iterations during training. The dark blue line represents a smoothed version of the training of the CNN, while the light blue line represents the accrual result from the training. The black striped line represents the accuracy results from the validation after 20 iterations.

**Figure 10.**
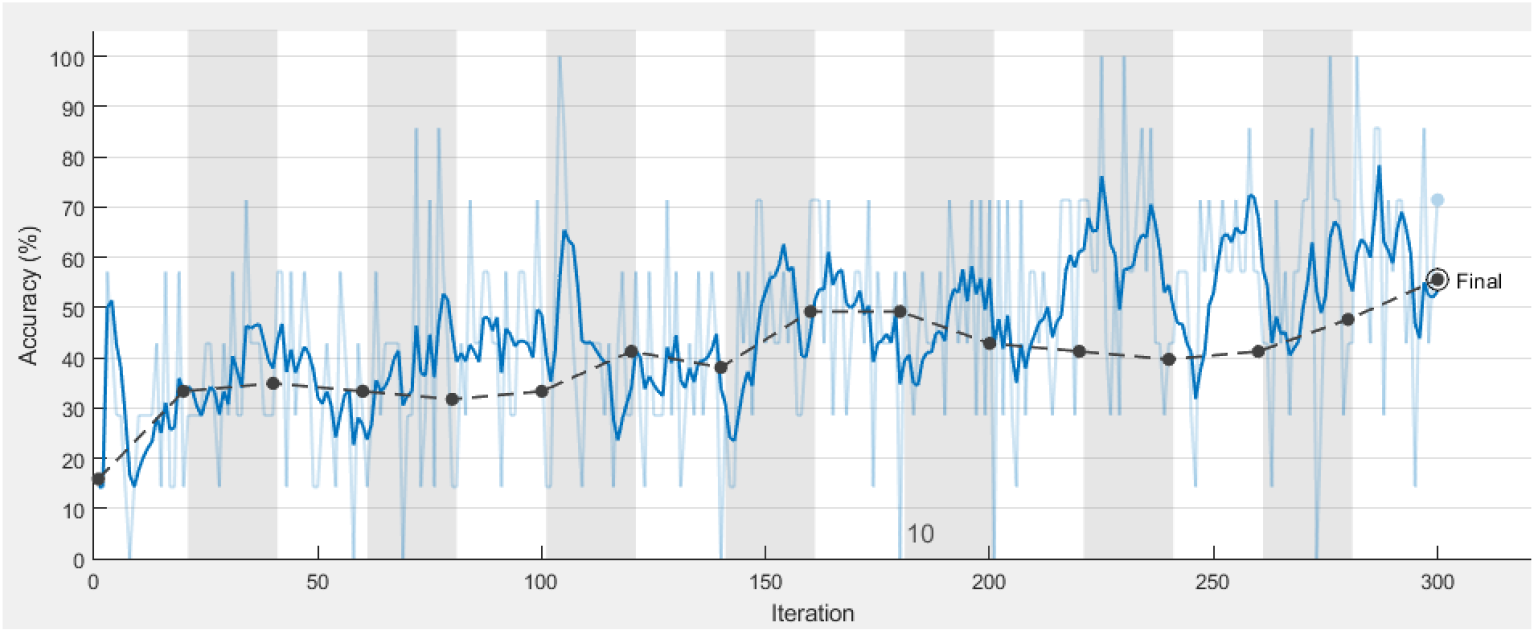
The accuracy of the Googlenet CNN over the iterations during training. The dark blue line represents a smoothed version of the training of the CNN, while the light blue line represents the accrual result from the training. The black striped line represents the accuracy results from the validation after 20 iterations.

## Discussion

The aim of the present study was to investigate the use of thermographic imaging and microphones (sound) in obtaining objective indicators for acute stress in laying hens. The results showed a decrease in average surface temperature when the birds were stressed. This decrease was most apparent in the beak and comb region. Various research has looked at the use of thermography in heat stress (Giloh et al., 2012; Marelli et al., 2012; Kim et al., 2021). Less research has been done on acute behavioural stress. One study found a decrease in temperature in the comb region after the birds were restricted (Herborn et al., 2015). Meanwhile, in this study, the eye and the head region also decreased in temperature. This decrease was still within the margin of error of the thermographic camera (± 1°C). Thus, this research can not decisively conclude that a decrease in temperature is occurring in these regions.

The beak of a bird has an important thermoregulatory function. Research in several tropical birds has shown that the larger beak these animals possess serves to cool the body down in higher temperatures (Ven et al., 2016). Like these tropical birds, the beak of a chicken is also highly vascularized and thus most likely serves a thermoregulatory function (Iqbal and Moss, 2021). From the results of the present study, it can be concluded that function extends to stress-induced thermoregulatory changes.

During acute stress the activation of the HPA-axis causes animals’ core temperature to rise (Oka, 2018). This is done by increasing vasodilation in the muscles and vasoconstriction in the skin’s blood vessels and peripheral regions such as digits. The allocation of more blood away from the skin increases muscle function in time of distress. From the results of the present study, it can be concluded that this reaction also takes place in younger chicks. The metabolic rate in older birds is higher, thus they will produce more heat. This makes a direct comparison between chicks and fully grown adult unreliable. Also, the stressor used in this study has not been used in conventional research in poultry stress. The pro of using this acute stressor over for example the tonic immobility test is that the chicken does not have to be separated from its pen mates. However, the con is that taking controlled measurements is more difficult, since the birds move around.

A recent study found similar results in the beak when the birds were exposed to tonic immobility (Soroko and Zaborski, 2021). The decrease observed in the present study was higher than the increase seen in the study by Soroko and Zaborski. This can be attributed to several reasons. The birds used in the study by Soroka and Zaborski used birds tat were 396 days old. Many environmental and physiological factors influence an animal’s thermoregulation, like the age, sex, ambient temperature and humidity (Oka, 2018). The way the regions of interest were drawn differed from the study by Soroko and Zaborski. A temperature difference was observed between the nostril and the beak. Both regions were included in the beak area for this research. Furthermore, the ambient temperature during the current study changed over time as the birds aged. The results show that the ambient temperature had a significant correlation with the temperature of the regions of interest.

Although the thermographic imaging showed potential technique for contactless monitoring of stress there are number of limitations should be addressed here. The angle at which a thermal image is taken relative to the target has an influence on the results that are collected (Tabh et al., 2021). During the current research, the birds were not restricted in their movement since this would have influenced the stress levels in the birds. The effect of this was that the birds were not always at the same distance or angle as the camera. Whilst the results were significant, large variations were found within the data. This can be attributed to a difference in the metabolism of the birds, a process already observed in mice (Lecorps et al., 2016). A major factor in this variability comes from the individual energy expenditure during physical activity. More active animals will produce a higher basal core temperature and this will influence their thermal stress response (Lecorps et al., 2016). Furthermore, individual chickens might react differently when exposed to a stressor, based on how they perceive the threat (Travain and Valsecchi, 2021). Further research should include some kind of identification to control for these individual differences.

The current price of the infrared technology used in this study is high. Research has shown that there is definitely a difference in readings between the different cameras (Cândido et al., 2018). However, if appropriate corrections were made to the data from the lower quality camera, it could be used as well as the higher quality camera. Although, this correction could not always be used. At higher skin temperatures (35°C), the low quality camera could not provide adequate results (Cândido et al., 2018). The problem with lower quality is also the subtlety that is lost with a lower resolution. In the present study, the eye was often small due to the size of the younger birds and the distance to the object. If this accuracy is lost, it could mean that influence from the body temperature, which is significantly higher than the eye temperature, could interfere with the true temperature in the smaller body parts. As was seen in the present study, the different areas of the body have a differing intensity in the reaction to stress. It is important to focus on these body parts to get an accurate assessment of the animals’ thermogenic reactions to stress.

Chickens would become unrestful from exposure to stress (Curtin et al., 2014). This unrestful state would have resulted in more vocalisations produced by the birds. In the present study, during the recording, the chickens did respond to the opening of the umbrella audibly, and this did not represent itself in the results. It could be a possibility that after the initial flight response described by Curtin et al. 2014, the chickens go into a state of more vigilant behaviour. This can result in so-called freezing behaviour (Amorim and Dias, 2021). However, freezing behaviour is often categorised as a short interruption of movement followed by unrest. Thus, it is unsure whether the decrease in calls resulted from this. Another reason for the decrease in calls might be other behaviours performed by the chicks. Since the prestress and post-stress recording were always taken at the same time of day, the daily behaviours of the birds could have had an influence. Birds might perform certain behaviours at certain times. These behaviours could correlate with various vocalisations and thus change the intensity and amount of calls. Other research on birds has shown that the amount of calls songbirds produce will fluctuate over the day (Mishra et al., 2020). Since the time of measuring the prestress situation was in the morning, it could also be concluded that the birds were more active during this time and thus produced more sounds. This could mean that the stress event had less of an influence on the number of calls. However, since a significant interaction between the exposure to stress and the treatment was found it could be a combination of the daily rhythm and the effect of acute stress that dictates the number of vocalisations. Future study could focus on mapping the daily behaviours related to their calls for chickens specifically.

During the observations, the birds in the different cages could not be separated from the other cages with a soundproof barrier. Thus, it cannot be excluded that some calls of the birds from the treatment cages appeared on the control audio file and vice versa. The MATLAB algorithm created for this research did not work properly for noisy environments. At times where the moisturizer or heaters were on the sound from the machines was overlapping with the calls from the chicks. While techniques exist that can filter these sounds it was out of the scope for this research to implement. This is also the case for neural network analysis. While the categorisation was able to be done at an 86 percent accuracy, it is unsure whether this comes from the stress condition of the birds or because of the proximity of the microphones to certain machinery. The points previously mentioned like the daily behavioural rhythm could also have influenced this method.

The use of audio measurements in birds does show a potential in detecting stress responses. Research has been done on small groups of birds, but more research should be done on large flocks to see what the challenges are in implementing audio measurement on a larger scale. The use of machine learning showed promise as a potential tool in this regard. Automatic recognition of specific calls could give great insight into chickens’ behaviour and physiological state. Future studies are warranted to further investigate the classification of vocalizations datasets from laying hens for complete validation. These datasets could be comprised of laying hens or broilers with health challenges along with control healthy bird’s audio signals.

## Conclusions

This research aimed to quantify the response to an acute stress factor in laying hens using two different kinds of sensors, namely, thermographic imagery and microphones. The results have shown that the skin temperature decreased significantly shortly after acute stress. This was most apparent in the beak and comb regions. Post-stress it was found that the number of vocalisations the birds produced decreased. This could be attributed to a freezing response or because of daily behavioural rhythm of the birds. Chickens might be more active during certain parts of the day and thus produce more sounds. In this study, the application of deep learning methods for automated laying hen’s vocalization classification was investigated. The results from the experiments of our study demonstrate that an ensemble of deep learning-based CNN approaches achieved a maximum accuracy of 85.7% in the laying hen vocalization classification. The results and methods for both thermographic imaging and audio analysis emphasise the importance of more research into the validation of these sensors with different ages, environmental conditions, and other stressors.

## Acknowledgements

The authors thank the Chair of Adaptation Physiology Group Professor Dr. Bas Kemp for his support of this study. The authors also thank the technicians of the animal care facility CARUS who helped us in the daily management of the birds and assistance during data collection. The authors express sincere thanks to Dr. Aaart Lammers for the valuable discussions and support in the setting up of animal experiments and for his help with the approval for animal care and experimental approval from the Wageningen University’ s Animal Experiments Board and IvD.

## Appendix

### Appendix I

Daily Maximum Air Temperature

**Table 3.** The maximum daily air temperature (°C) for the control cage (1) and the treatment cages (2)

| Date | Age of birds (days) | Cage | Max Temperature (°C) |
| --- | --- | --- | --- |
| 16-nov | 1 | 1 | 33.4 |
| 16-nov | 1 | 2 | 34.9 |
| 17-nov | 2 | 1 | 34 |
| 17-nov | 2 | 2 | 34.5 |
| 18-nov | 3 | 1 | 34.2 |
| 18-nov | 3 | 2 | 34.5 |
| 19-nov | 4 | 1 | 34 |
| 19-nov | 4 | 2 | 34 |
| 20-nov | 5 | 1 | 33 |
| 20-nov | 5 | 2 | 33 |
| 21-nov | 6 | 1 | 32.6 |
| 21-nov | 6 | 2 | 32.1 |
| 22-nov | 7 | 1 | 32.2 |
| 22-nov | 7 | 2 | 32.5 |
| 23-nov | 8 | 1 | 31.5 |
| 23-nov | 8 | 2 | 31.8 |
| 24-nov | 9 | 1 | 31 |
| 24-nov | 9 | 2 | 31.2 |
| 25-nov | 10 | 1 | 30.8 |
| 25-nov | 10 | 2 | 31 |
| 26-nov | 11 | 1 | 30.5 |
| 26-nov | 11 | 2 | 30.9 |
| 27-nov | 12 | 1 | 31.7 |
| 27-nov | 12 | 2 | 30 |
| 28-nov | 13 | 1 | 30.5 |
| 28-nov | 13 | 2 | 28.4 |
| 29-nov | 14 | 1 | 30.5 |
| 29-nov | 14 | 2 | 28.4 |
| 30-nov | 15 | 1 | 31.3 |
| 30-nov | 15 | 2 | 30.2 |
| 1-dec | 16 | 1 | 31.4 |
| 1-dec | 16 | 2 | 30.5 |
| 2-dec | 17 | 1 | 31.3 |
| 2-dec | 17 | 2 | 30.3 |
| 3-dec | 18 | 1 | 28.5 |
| 3-dec | 18 | 2 | 28.8 |
| 4-dec | 19 | 1 | 27.9 |
| 4-dec | 19 | 2 | 27.9 |
| 5-dec | 20 | 1 | 27.2 |
| 5-dec | 20 | 2 | 27.3 |
| 6-dec | 21 | 1 | 27.5 |
| 6-dec | 21 | 2 | 27.5 |
| 7-dec | 22 | 1 | 28 |
| 7-dec | 22 | 2 | 28.7 |
| 8-dec | 23 | 1 | 27.2 |
| 8-dec | 23 | 2 | 27.8 |
| 9-dec | 24 | 1 | 26.6 |
| 9-dec | 24 | 2 | 27.2 |
| 10-dec | 25 | 1 | 27 |
| 10-dec | 25 | 2 | 27.2 |
| 11-dec | 26 | 1 | 28.9 |
| 11-dec | 26 | 2 | 27.5 |
| 12-dec | 27 | 1 | 27.6 |
| 12-dec | 27 | 2 | 26 |
| 13-dec | 28 | 1 | 28.2 |
| 13-dec | 28 | 2 | 26.3 |
| 14-dec | 29 | 1 | 27.9 |
| 14-dec | 29 | 2 | 26.5 |
| 15-dec | 30 | 1 | 27.9 |
| 15-dec | 30 | 2 | 26.4 |
| 16-dec | 31 | 1 | 25.2 |
| 16-dec | 31 | 2 | 24.9 |
| 17-dec | 32 | 1 | 25.8 |
| 17-dec | 32 | 2 | 28.5 |
| 18-dec | 33 | 1 | 25.4 |
| 18-dec | 33 | 2 | 26 |
| 19-dec | 34 | 1 | 25.2 |
| 19-dec | 34 | 2 | 24.8 |
| 20-dec | 35 | 1 | 24.9 |
| 20-dec | 35 | 2 | 24 |
| 21-dec | 36 | 1 | 24.8 |
| 21-dec | 36 | 2 | 25.1 |
| 22-dec | 37 | 1 | 23.1 |
| 22-dec | 37 | 2 | 26.2 |
| 23-dec | 38 | 1 | 23.9 |
| 23-dec | 38 | 2 | 25.3 |
| 24-dec | 39 | 1 | 23.8 |
| 24-dec | 39 | 2 | 25.1 |
| 25-dec | 40 | 1 | 23.6 |
| 25-dec | 40 | 2 | 24.8 |
| 26-dec | 41 | 1 | 23.8 |
| 26-dec | 41 | 2 | 25.4 |
| 27-dec | 42 | 1 | 23 |
| 27-dec | 42 | 2 | 25.7 |
| 28-dec | 43 | 1 | 23.4 |
| 28-dec | 43 | 2 | 25.3 |
| 29-dec | 44 | 1 | 23.7 |
| 29-dec | 44 | 2 | 24.4 |
| 30-dec | 45 | 1 | 24 |
| 30-dec | 45 | 2 | 24.5 |
| 31-dec | 46 | 1 | 24.5 |
| 31-dec | 46 | 2 | 24.8 |
| 1-jan | 47 | 1 | 24.2 |
| 1-jan | 47 | 2 | 24.9 |
| 2-jan | 48 | 1 | 23.7 |
| 2-jan | 48 | 2 | 24.7 |
| 3-jan | 49 | 1 | 24 |
| 3-jan | 49 | 2 | 24.9 |
| 4-jan | 50 | 1 | 23.7 |
| 4-jan | 50 | 2 | 24.3 |
| 5-jan | 51 | 1 | 22.9 |
| 5-jan | 51 | 2 | 23.8 |
| 6-jan | 52 | 1 | 22.4 |
| 6-jan | 52 | 2 | 23.6 |
| 7-jan | 53 | 1 | 22.6 |
| 7-jan | 53 | 2 | 22 |
| 8-jan | 54 | 1 | 22.9 |
| 8-jan | 54 | 2 | 22.1 |
| 9-jan | 55 | 1 | 22.8 |
| 9-jan | 55 | 2 | 23.7 |
| 10-jan | 56 | 1 | 22.8 |
| 10-jan | 56 | 2 | 23.7 |
| 11-jan | 57 | 1 | 22.3 |
| 11-jan | 57 | 2 | 23 |
| 12-jan | 58 | 1 | 22 |
| 12-jan | 58 | 2 | 22 |
| 13-jan | 59 | 1 | 22 |
| 13-jan | 59 | 2 | 22.7 |
| 14-jan | 60 | 1 | 22.2 |
| 14-jan | 60 | 2 | 22.8 |
| 15-jan | 61 | 1 | 22.2 |
| 15-jan | 61 | 2 | 21.7 |
| 16-jan | 62 | 1 | 22 |
| 16-jan | 62 | 2 | 21.8 |
| 17-jan | 63 | 1 | 22.5 |
| 17-jan | 63 | 2 | 22.7 |
| 18-jan | 64 | 1 | 22.2 |
| 18-jan | 64 | 2 | 22.8 |

### Appendix II

Data Collection Period / Schedule

**Table 4.**
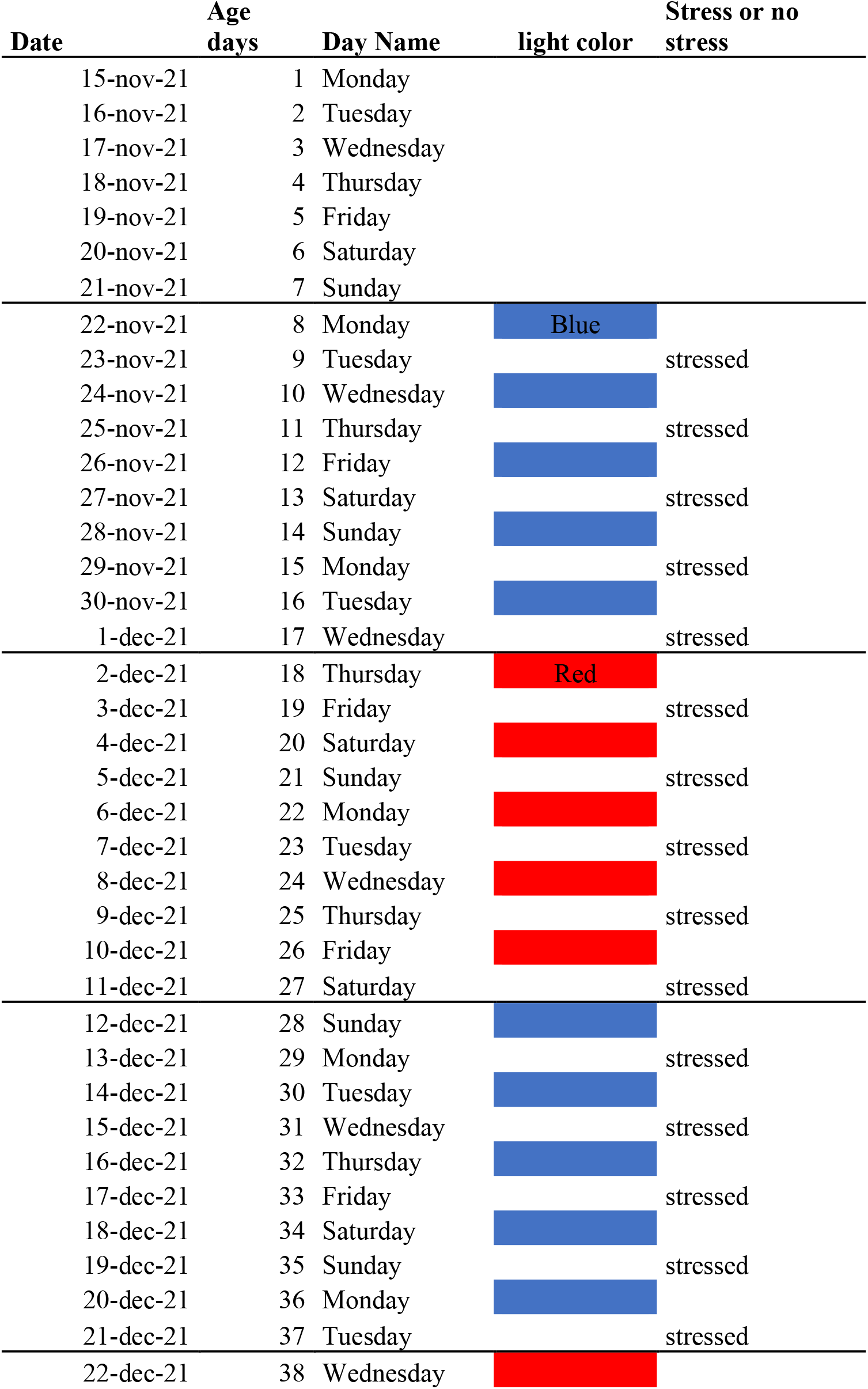
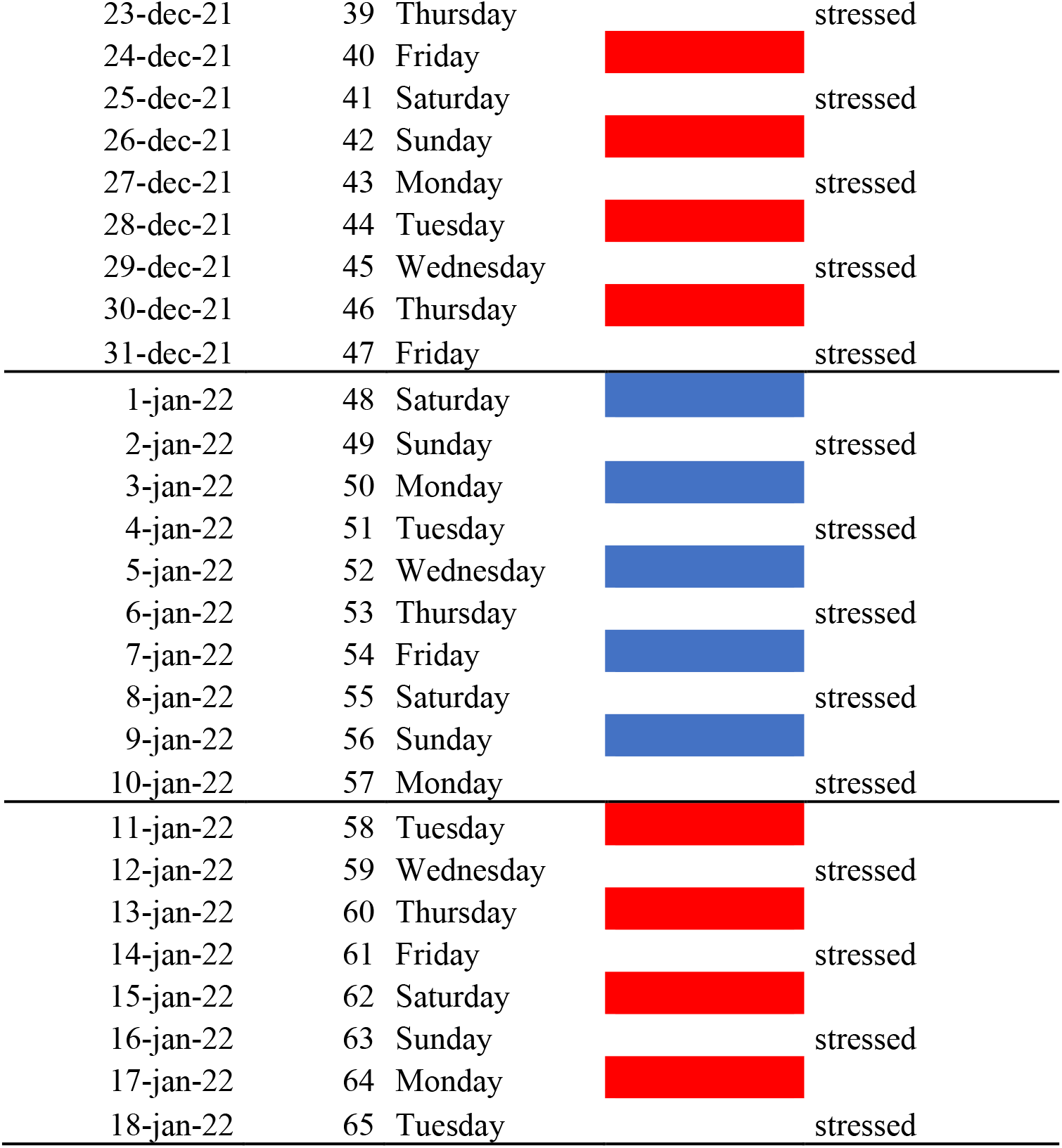
The lighting and stressing schedule for the study. The chickens were also part of another study that researched the effects of different light colours

| Date | Age days | Day Name | light color | Stress or no stress |
| --- | --- | --- | --- | --- |
| 15-nov-21 | 1 | Monday |  |  |
| 16-nov-21 | 2 | Tuesday |  |  |
| 17-nov-21 | 3 | Wednesday |  |  |
| 18-nov-21 | 4 | Thursday |  |  |
| 19-nov-21 | 5 | Friday |  |  |
| 20-nov-21 | 6 | Saturday |  |  |
| 21-nov-21 | 7 | Sunday |  |  |
| 22-nov-21 | 8 | Monday | Blue |  |
| 23-nov-21 | 9 | Tuesday |  | stressed |
| 24-nov-21 | 10 | Wednesday |  |  |
| 25-nov-21 | 11 | Thursday |  | stressed |
| 26-nov-21 | 12 | Friday |  |  |
| 27-nov-21 | 13 | Saturday |  | stressed |
| 28-nov-21 | 14 | Sunday |  |  |
| 29-nov-21 | 15 | Monday |  | stressed |
| 30-nov-21 | 16 | Tuesday |  |  |
| 1-dec-21 | 17 | Wednesday |  | stressed |
| 2-dec-21 | 18 | Thursday | Red |  |
| 3-dec-21 | 19 | Friday |  | stressed |
| 4-dec-21 | 20 | Saturday |  |  |
| 5-dec-21 | 21 | Sunday |  | stressed |
| 6-dec-21 | 22 | Monday |  |  |
| 7-dec-21 | 23 | Tuesday |  | stressed |
| 8-dec-21 | 24 | Wednesday |  |  |
| 9-dec-21 | 25 | Thursday |  | stressed |
| 10-dec-21 | 26 | Friday |  |  |
| 11-dec-21 | 27 | Saturday |  | stressed |
| 12-dec-21 | 28 | Sunday |  |  |
| 13-dec-21 | 29 | Monday |  | stressed |
| 14-dec-21 | 30 | Tuesday |  |  |
| 15-dec-21 | 31 | Wednesday |  | stressed |
| 16-dec-21 | 32 | Thursday |  |  |
| 17-dec-21 | 33 | Friday |  | stressed |
| 18-dec-21 | 34 | Saturday |  |  |
| 19-dec-21 | 35 | Sunday |  | stressed |
| 20-dec-21 | 36 | Monday |  |  |
| 21-dec-21 | 37 | Tuesday |  | stressed |
| 22-dec-21 | 38 | Wednesday |  |  |
| 23-dec-21 | 39 | Thursday |  | stressed |
| 24-dec-21 | 40 | Friday |  |  |
| 25-dec-21 | 41 | Saturday |  | stressed |
| 26-dec-21 | 42 | Sunday |  |  |
| 27-dec-21 | 43 | Monday |  | stressed |
| 28-dec-21 | 44 | Tuesday |  |  |
| 29-dec-21 | 45 | Wednesday |  | stressed |
| 30-dec-21 | 46 | Thursday |  |  |
| 31-dec-21 | 47 | Friday |  | stressed |
| 1-jan-22 | 48 | Saturday |  |  |
| 2-jan-22 | 49 | Sunday |  | stressed |
| 3-jan-22 | 50 | Monday |  |  |
| 4-jan-22 | 51 | Tuesday |  | stressed |
| 5-jan-22 | 52 | Wednesday |  |  |
| 6-jan-22 | 53 | Thursday |  | stressed |
| 7-jan-22 | 54 | Friday |  |  |
| 8-jan-22 | 55 | Saturday |  | stressed |
| 9-jan-22 | 56 | Sunday |  |  |
| 10-jan-22 | 57 | Monday |  | stressed |
| 11-jan-22 | 58 | Tuesday |  |  |
| 12-jan-22 | 59 | Wednesday |  | stressed |
| 13-jan-22 | 60 | Thursday |  |  |
| 14-jan-22 | 61 | Friday |  | stressed |
| 15-jan-22 | 62 | Saturday |  |  |
| 16-jan-22 | 63 | Sunday |  | stressed |
| 17-jan-22 | 64 | Monday |  |  |
| 18-jan-22 | 65 | Tuesday |  | stressed |

### Appendix III

Confusion Matrix for Convolutional Neural Network Analysis of Vocalizations Data

**Table 5.**
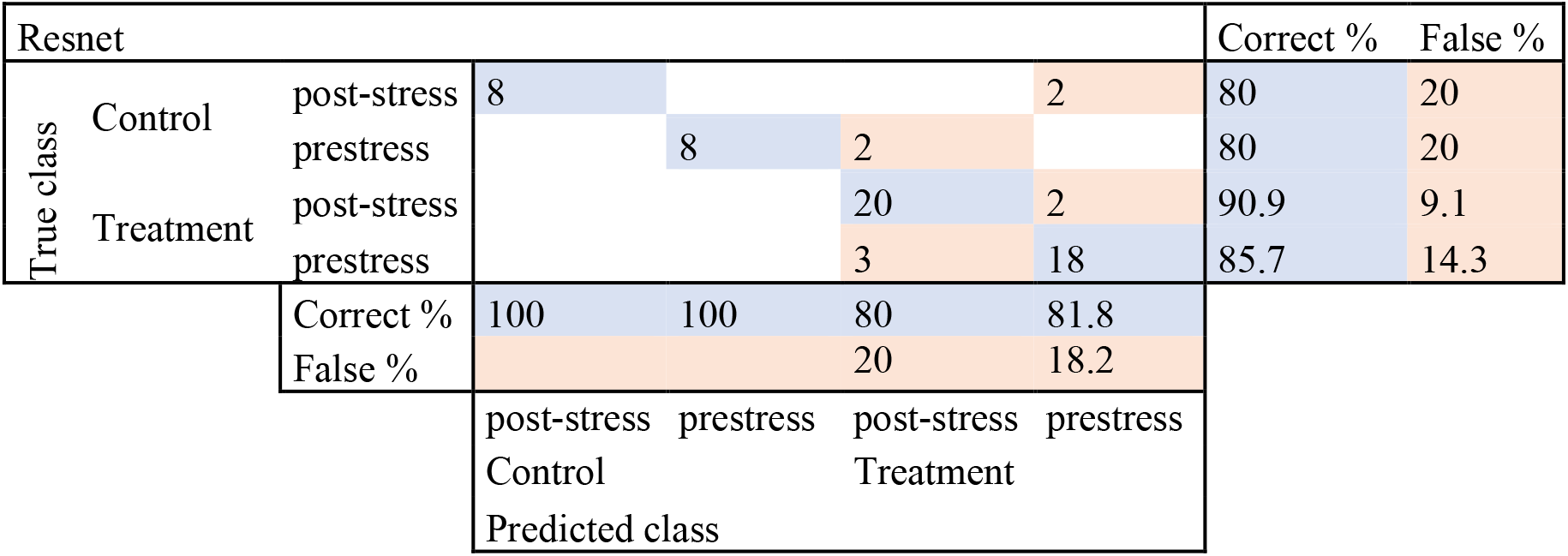
The classification accuracy for the Resnet method showing where the different files were categorised by the neural network. The true class is the actual class a file belongs to, and the predicted class is the place where the algorithm placed the file. The blue cells represent a correct classification, and the orange cells represent an incorrect classification

**Table 6.**
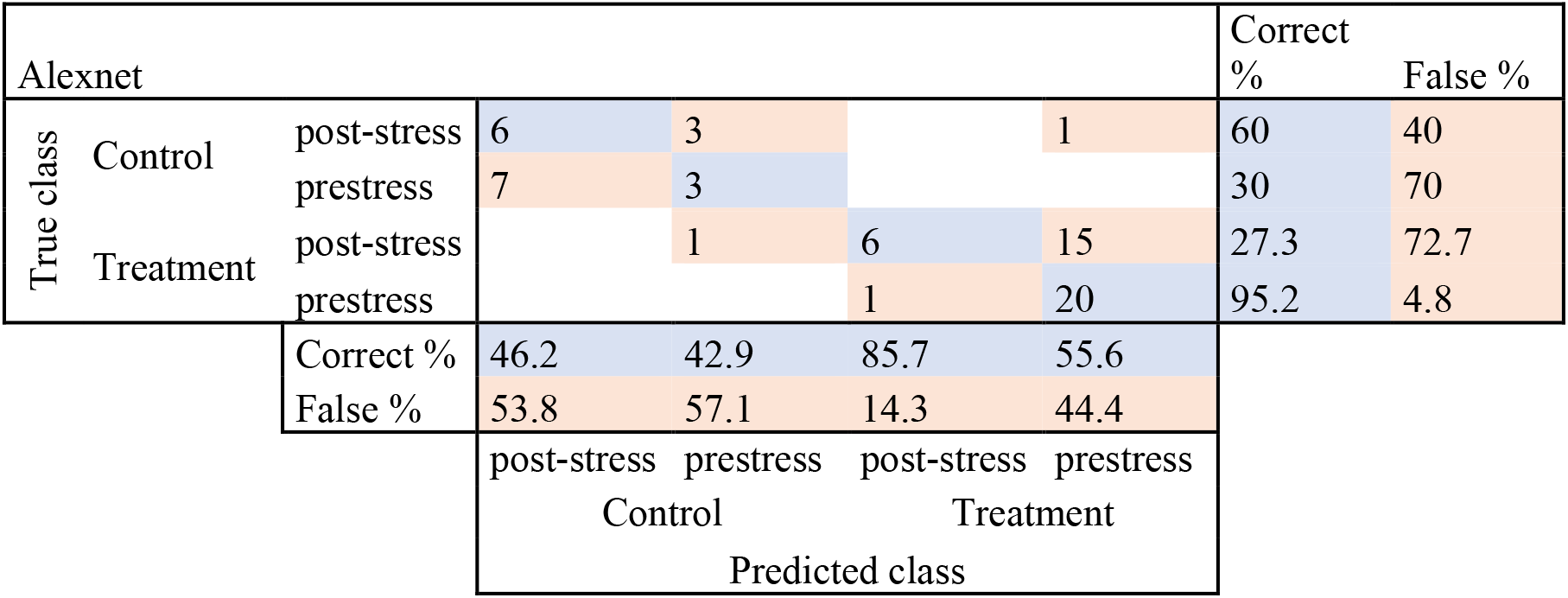
The classification accuracy for the Alexnet method showing where the different files were categorised by the neural network. The true class is the actual class a file belongs to and the predicted class is the place where the algorithm placed the file. The blue cells represent a correct classification and the orange cells represent an incorrect classification

**Table 7.**
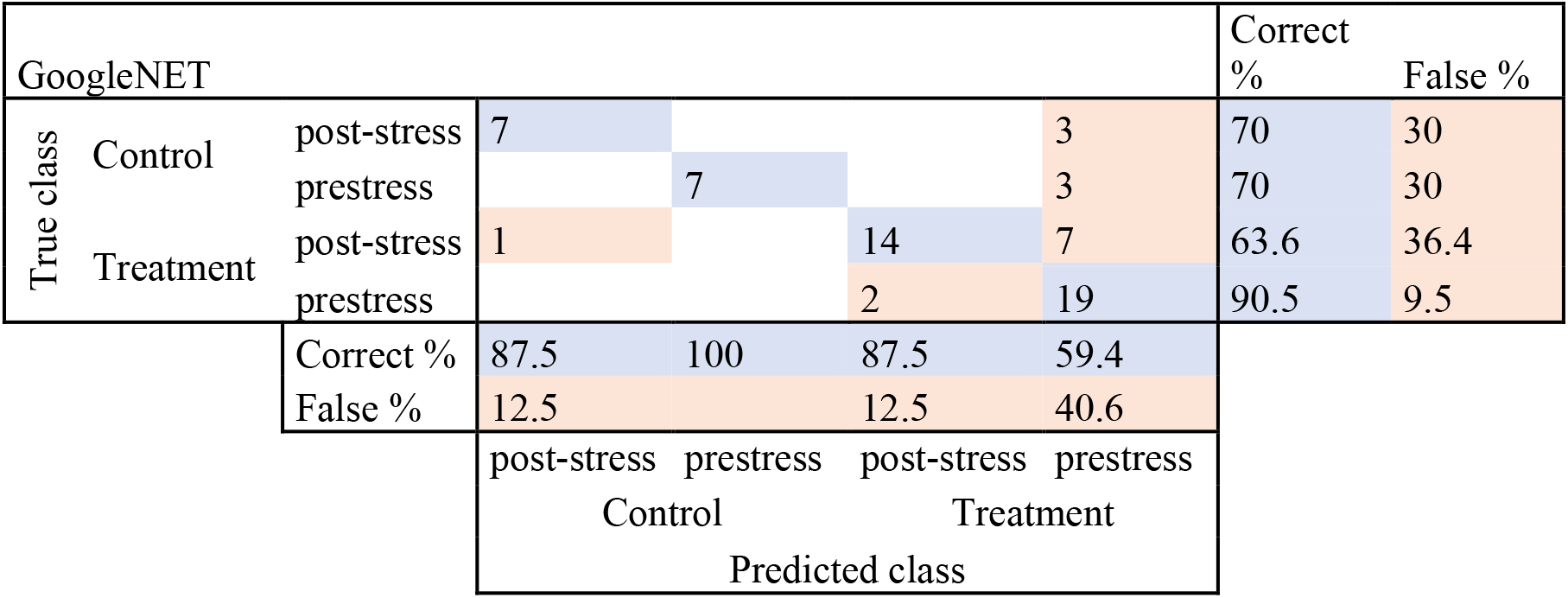
The classification accuracy for the GoogleNET method showing where the different files were categorised by the neural network. The true class is the actual class a file belongs to, and the predicted class is the place where the algorithm placed the file. The blue cells represent a correct classification, and the orange cells represent an incorrect classification.

## Notes

### Competing Interest Statement

The authors have declared no competing interest.

